# Genetic drift dominates genome-wide regulatory evolution following an ancient whole genome duplication in Atlantic salmon

**DOI:** 10.1101/2020.11.20.389684

**Authors:** Jukka-Pekka Verta, Henry Barton, Victoria Pritchard, Craig Primmer

## Abstract

Whole genome duplications (WGD) have been considered as springboards that potentiate lineage diversification through increasing functional redundancy. Divergence in gene regulatory elements is a central mechanism for evolutionary diversification, yet the patterns and processes governing regulatory divergence following events that lead to massive functional redundancy, such as WGD, remain largely unknown. We studied the patterns of divergence and strength of natural selection on regulatory elements in the Atlantic salmon (*Salmo salar*) genome, which has undergone WGD 100-80 Mya. Using ChIPmentation, we first show that H3K27ac, a histone modification typical to enhancers and promoters, is associated with genic regions, tissue specific transcription factor binding motifs, and with gene transcription levels in immature testes. Divergence in transcription between duplicated genes from WGD (ohnologs) correlated with difference in the number of proximal regulatory elements, but not with promoter elements, suggesting that functional divergence between ohnologs after WGD is mainly driven by enhancers. By comparing H3K27ac regions between duplicated genome blocks, we further show that a longer polyploid state post-WGD has constrained asymmetric regulatory evolution. Patterns of genetic diversity across natural populations inferred from re-sequencing indicate that recent evolutionary pressures on H3K27ac regions are dominated by largely neutral evolution. In sum, our results suggest that post-WGD functional redundancy in regulatory elements continues to have an impact on the evolution of the salmon genome, promoting largely neutral evolution of regulatory elements despite their association with transcription levels. These results highlight a case where genome-wide regulatory evolution following an ancient WGD is dominated by genetic drift.

**Significance statement:** Regulatory evolution following whole genome duplications (WGD) has been investigated at the gene expression level, but studies of the regulatory elements that control expression have been lacking. By investigating regulatory elements in the Atlantic salmon genome, which has undergone a whole genome duplication 100-80 million years ago, we discovered patterns suggesting that neutral divergence is the prevalent mode of regulatory element evolution post-WGD. Our results suggest mechanisms for explaining the prevalence of asymmetric gene expression evolution following whole genome duplication, as well as the mismatch between evolutionary rates in enhancers versus that of promoters.

## Introduction

Numerous evolutionary innovations have taken place in conjunction with, or following, large-scale genomic rearrangements such as whole genome duplications (WGD) (Vandepoele et al. 2004; Peer et al. 2009; Macqueen and Johnston 2014), which create massive functional redundancy of genes and regulatory elements. The effects of increased functional redundancy are thought to reflect onto gene expression evolution, albeit being varied and contingent on the function of the genes impacted (Robertson et al. 2017; Hallin and Landry 2019). Yet, the impacts of whole genome duplications on patterns of regulatory divergence and evolutionary forces acting on regulatory elements remain poorly documented and understood.

Gene expression levels and patterns in metazoan genomes are controlled by enhancer and promoter sequences commonly referred to as regulatory elements (Long et al. 2016). Regulatory elements contain recognition motifs for transcription factors (TFs) that cooperatively orchestrate gene expression of their target genes. Regulation of mammalian gene expression is putatively controlled by hundreds of thousands of regulatory elements (Villar et al. 2014; Gasperini et al. 2020), their number greatly exceeding the typical number of genes. Through their combined and individual function, regulatory elements have long been hypothesized to significantly contribute to evolutionary change and adaptation (King and Wilson 1975; Stern and Orgogozo 2008).

Gene regulatory elements can be mapped genome-wide by their distinct chromatin structure without knowledge of the specific TFs that recognize each element (Andersson and Sandelin 2019). Chromatin immunoprecipitation followed by high through-put sequencing (ChIP-seq) of H3K27 acetylation (H3K27ac), an epigenetic mark associated with active enhancers and promoters (Creyghton et al. 2010), has been particularly informative for studies on the evolution of regulatory elements because of its high phylogenetic conservation. Widely applicable ChIP-seq strategies for H3K27ac have facilitated studies on the evolution of regulatory elements across species, revealing that enhancers in particular can evolve rapidly (Villar et al. 2015). Past focus on regulatory element evolution has been dominated by studies in mammals (Villar et al. 2015; Berthelot et al. 2018), with little information on regulatory evolution in species that have more complex genomes due to more or less ancient WGD (Elurbe et al. 2017). Studying regulatory element evolution in post-WGD genomes is essential for a more comprehensive understanding how functional redundancy impacts regulatory evolution and species diversity.

Salmonid fishes are a promising system for studying regulatory evolution in post-WGD genomes. Salmonids underwent a WGD 100-80 million years ago (Mya) that resulted in a portion of the genome remaining in effective polyploidy (Lien et al. 2016). Despite massive gene and functional redundancy still existing in salmonid genomes, most gene duplicates from WGD (hereafter ohnologs, called homeologs in (Lien et al. 2016) or WGD paralogs) can be confidently identified, facilitating broad-scale genomic studies of ohnolog duplicate pairs. Across salmonids, gene expression evolution is dominated by asymmetric evolution where ohnologs retain and lose expression patterns in an unbalanced pattern (Lien et al. 2016; Gillard et al. 2020). Asymmetric expression evolution of ohnologs suggests that expression changes are driven by divergence in *cis-*acting regulatory elements such as enhancers and promoters because both gene copies are exposed to the same *trans*-acting factors within a cell. Studying evolutionary dynamics in gene regulatory regions however requires a map of enhancer and promoter elements.

Here, we studied gene regulatory elements in Atlantic salmon (*Salmo salar*), as species with a relatively recent WGD. By mapping regulatory elements in the tissue with the most transcribed genes, the testis, we characterized the association of H3K27ac with gene transcription and the presence of transcription factor recognition motifs. We compared regulatory elements between ohnologs and large-scale duplicate regions called homeoblocks, and discovered patterns consistent with large-scale, neutral divergence of regulatory elements. We further modelled the current strength of selection on regulatory elements using population re-sequencing data to show that regulatory elements continue to experience predominantly neutral evolution. Our results point towards a patterns of largely neutral regulatory element evolution within this post-WGD genome.

## Results

### H3K27ac ChIPmentation peaks associate with gene regions

To address the paucity of regulatory element genetic maps in species with recently duplicated genomes, we used ChIPmentation (Schmidl et al. 2015) of H3K27ac in immature salmon testis to map putative regulatory regions of the Atlantic salmon genome. Immature testes express the broadest number of salmon transcripts among 14 tested tissues (Fig S1), making testis well-suited for inferring regulatory regions in a genome-wide manner using a single tissue. Our analysis detected 34,489 reproducible H3K27ac regions (hereafter “peaks”, FDR<0.05 & logFC>1, N=5, Fig 1A). H3K27ac peak widths formed a marked periodical pattern with a median size of 623 bp, supporting the peaks corresponding to sets of nucleosomes with acetylated histones (Fig S2).

**Figure 1.**
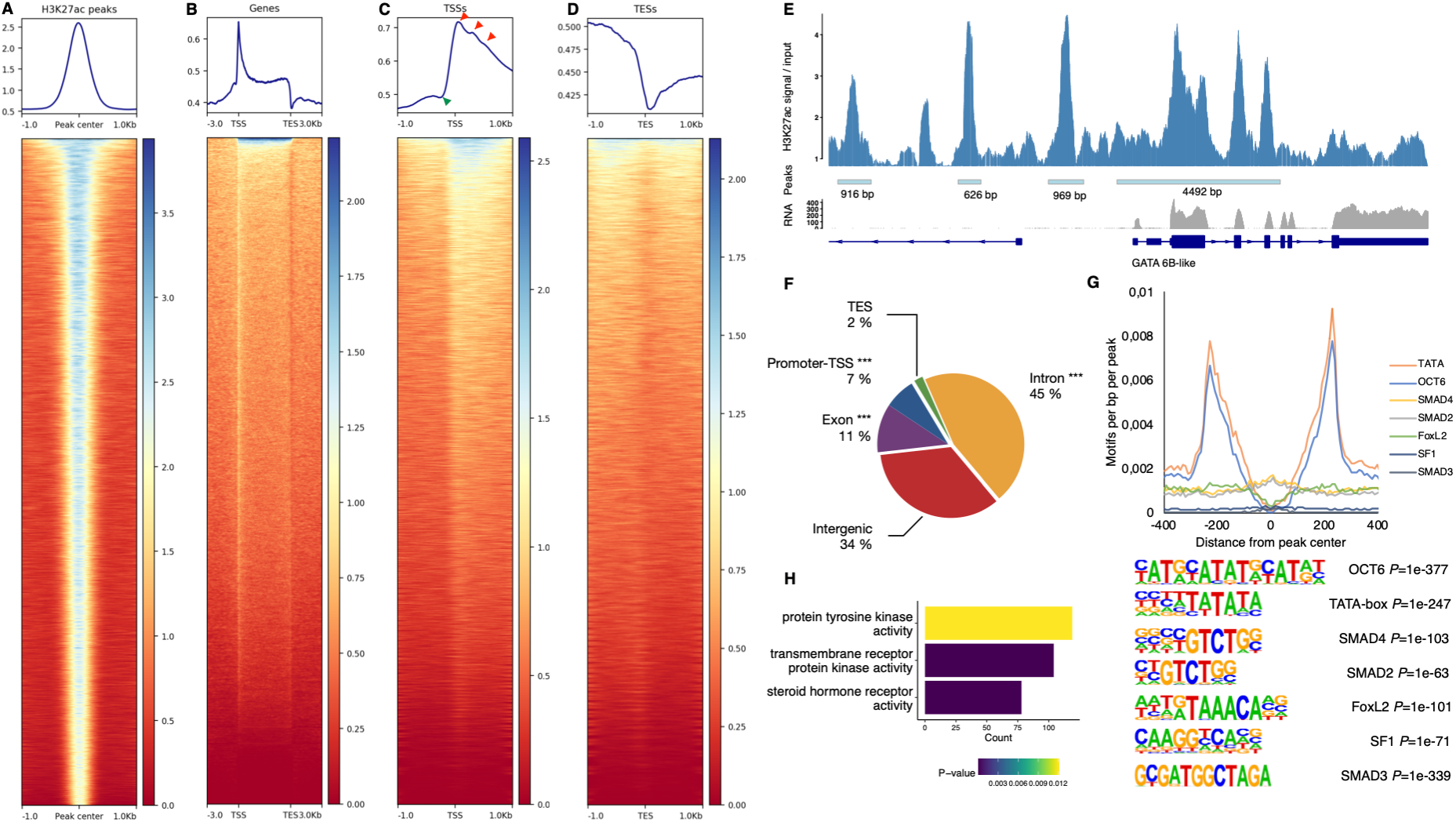
Key features of Atlantic salmon H3K27ac regions and associated genes in testis. (A) H3K27ac ChIPmentation signal over replicated H3K27ac peaks. (B) The majority of salmon genes show increased H3K27ac signal. (C) Transcription start sites (TSS) show particularly high H3K27ac signal with a prominent nucleosome-free region (NFR) immediately upstream of TSS. Red arrows correspond to +1, +2 and +3 nucleosomes, green arrow corresponds to promoter NFR. (D) Transcription end sites (TES) are characterized by a prominent NFR. (E) A region with prominent H3K27ac peaks overlapping the GATA 6B-like locus. (F) Division of 34,489 reproducible H3K27ac peaks according to genomic feature type. ***, likelihood-ratio overrepresentation *p*<0.001. (G) Density of six testis cell-type associated and over-represented TF motifs relative to H3K27ac peak center (upper panel). Motif sequences of TF motifs with their over-representation *P*-values (lower panel). (H) Significantly over-represented gene-ontology functions in genes with at least one proximal H3K27ac peak.

Regions containing genes showed a distinctive pattern of high H3K27ac signal, consistent with a highly organized nucleosome occupancy (Mavrich et al. 2008; Jiang and Pugh 2009). Strikingly, such organized nucleosome occupancy was observed for the majority of salmon genes (Fig 1B), consistent with most genes being actively transcribed in the testis. Transcription start site (TSS) regions were characterized by a dip in H3K27ac signal immediately upstream of the TSS corresponding to the 5’ nucleosome free region (NFR) observed in expressed genes (Fig 1C green arrow). Nucleosome patterning with acetylated H3K27 could also be observed for the three nucleosomes downstream from the TSS, with strongest signal on the first (+1) nucleosome (Fig 1C red arrows). Transcription end sites (TES) were characterized by a 3’ NFR void of H3K27ac signal, corresponding to the region of transcription termination (Fig 1D). The largest single H3K27ac region, comprising multiple individual H3K27ac peaks, overlapped with GATA 6B-like gene locus (Fig 1E), which is expressed in Sertoli and germ cells, and plays a central and conserved role in regulating testis gene expression and cell differentiation (Viger et al. 2008).

Overall, the distribution of H3K27ac peaks showed association with promoters, exons (including the 5’UTR), and especially introns, which contained the largest percentage, 45%, of the peaks (Fig 1F). We further used a set of peaks with 500 bp width and centered to nucleosome free regions to identify over-represented motifs in H3K27ac peaks. These replicated peaks overlapping 83% of the original peak set were over-represented in motifs for general promoter features (TATA-box 26.8% of peaks), as well as for transcription factors implicated in the development of spermatogonia (OCT6 10.5% of peaks), Sertoli and Leydig cells (SMAD4 41% of peaks, SMAD2 38.4% of peaks, FoxL2 36.7% of peaks, SF1 9.3% of peaks and SMAD3 2.3% of peaks). Plotting motif density relative to peak center revealed that TATA, OCT6 and FoxL2 motifs tended to be situated on the opposite site of acetylated nucleosomes, while SMAD and SF1 motifs were centered on peaks (Fig 1G). We hypothesize that this pattern can arise from the latter two motifs residing primarily in proximity to regions with well-defined NFR such as TSSs. Genes proximal to at least one H3K27ac peak were enriched for three GO functions notably involved in cell signaling and differentiation, consistent with the tight regulation of cell cycle in immature testis cell types (Fig 1H). Together these data demonstrate that H3K27ac regions in Atlantic salmon testis bear the hallmarks of actively transcribed genes and tissue-specific regulation of cell type differentiation.

### Divergence in proximal peaks tracks expression divergence between homeologs

To further determine whether the H3K27ac peaks we discovered were associated with regulatory activity, and therefore represented true gene regulatory elements, we investigated peaks in conjunction with expression levels of genes in immature testis RNA-seq data. We hypothesized that if H3K27ac peaks functioned in an additive manner, the number of peaks assigned to genes should correlate with their expression levels. Indeed, the number of promoter, and total proximal peaks assigned to nearest TSSs correlated positively with immature testis gene expression levels (Spearman’s rho 0.1, linear regression *p*<2.2e-16), with the number of promoter peaks having a stronger effect on expression level compared to total proximal peaks (Fig 2A). These results support the H3K27ac regions being functional enhancer and promoter elements with an additive effect on expression levels of proximal genes.

**Figure 2.**
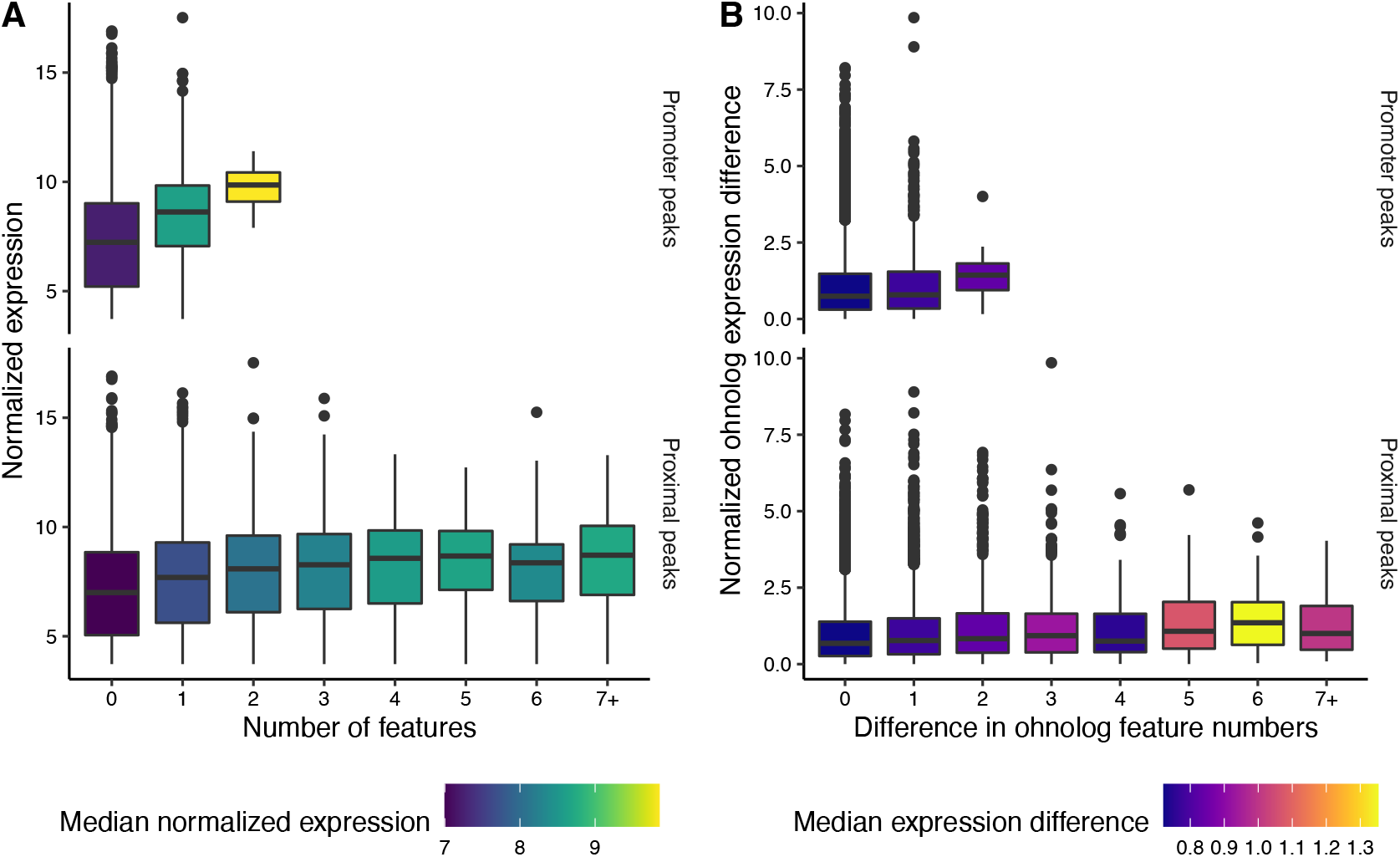
H3K27ac peaks are associated with gene expression levels. (A) Number of promoter and total proximal peaks correlates positively with gene expression levels in testis. (B) Difference in total proximal peaks, but not promoter peaks, correlates with expression differences between ohnologs from WGD.

We next investigated whether divergence in H3K27ac peaks between ohnologs from WGD had phenotypic impacts by comparing the number of assigned peaks and the expression levels between ohnologs. The number of peaks assigned to one ohnolog could not be used to predict the number of peaks assigned to the other (Fig S3), indicating a general absence of conservation for the number of regulatory elements for ohnologs. These results were robust to different strategies for identifying ohnologs (Supplementary analysis).

To test whether the observed differences in the number of peaks assigned to ohnologs had phenotypic effects, we compared the difference in peak number to expression differences between the ohnologs. Divergence in total proximal peak number was positively associated with divergence in mean expression levels (linear regression *p*<2.2e-16, Fig 2B), whereas the difference in only promoter peaks did not predict expression difference (linear regression *p*<0.072, Fig 2B). These results indicate that the number of peaks assigned to ohnologs seem to diverge in a random fashion, yet divergence in proximal H3K27ac peaks between ohnologs has a phenotypic impact of increasing the level of expression differences.

### Reploidization age of homeoblocks predicts the magnitude of divergence in the regulatory landscape

Whole genome duplications can leave large portions of the genome remaining in effective polyploidy until recent evolutionary history (Lien et al. 2016), yet the extent to which duplicated genome regions share, or differ in, regulatory elements today remains unknown. To investigate the large-scale trends in regulatory element evolution between duplicated genome regions from WGD, we first compared H3K27ac peaks assigned to homeoblocks. Overall, H3K27ac peak numbers were highly correlated between homeoblocks (linear regression R^2^=0.98, *p*<2.2e-16, Fig 3A). Normalising the number of H3K27ac peaks to the length of the homeoblocks revealed divergence from the 1:1 expected in H3K27ac peak density between blocks (Fig 3B). We hypothesised that if divergence in H3K27ac peak density between homeoblocks was a random process, we would expect to see that shorter homeoblocks have diverged more in peak density compared to longer homeoblocks. Difference from 1:1 peak density indeed followed block length in a manner suggesting a process of random peak loss/gain per unit of homeoblock length (Fig 3C).

**Figure 3.**
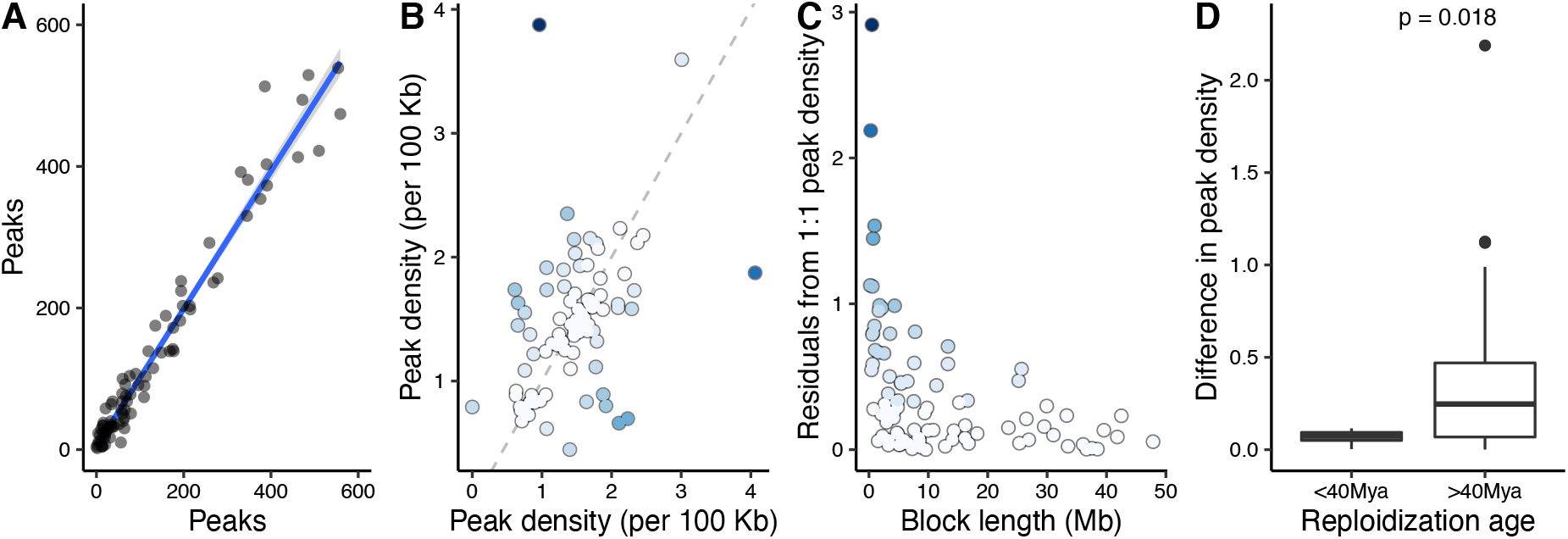
Comparison of H3K27ac peaks between homeoblocks from WGD. (A) The number of H3K27ac peaks is highly correlated between homeoblocks. (B) Peak density identifies homeoblocks that have diverged more from the expected 1:1 density (grey dashed line). Points are coloured for residuals from 1:1 line. (C) Peak density has diverged more in homeoblocks with shorter length. (D) Homeoblocks that have remained in effective polyploidy until more recently have diverged less in peak density.

The Atlantic salmon genome contains regions that have remained polyploid until as recently as 40 Mya, and which have been proposed to be implicated in salmonid adaptations (Robertson et al. 2017). Therefore, we tested whether the time since reploidization of homeoblocks influenced divergence in regulatory region density. Homeoblocks with reploidization age less than 40 Mya were significantly more conserved in H3K27ac peak density (*p*=0.018, Fig 3D), suggesting that a longer polyploid state of homeoblocks has constrained asymmetric regulatory divergence between the blocks.

### A genome-wide map of regulatory divergence in homeoblocks

We next investigated the genomic distribution of H3K27ac peaks to understand whether certain regions of the genome showed higher peak densities or differences therein. All chromosomes showed regulatory region activity (Fig 4), and overall, the number of peaks in 1 Mb bins along chromosomes was correlated to the number of expressed genes in the same bins (R^2^=0.08, linear regression *p*<2.2e-16, Fig S4). There were also indications that homeoblocks with most elevated differences in H3K27ac density per 100 Kb of block length tended to group to certain chromosomes. Notably, chromosome 19 contained three of the ten blocks with the most diverged peak density (hypergeometric test *p*<0.012), followed by chromosome 1 containing four (hypergeometric test *p*<0.018), and chromosome 9 containing three (hypergeometric test *p*<0.012) (each block is present in two chromosomes by definition). The ten most diverged homeoblocks accounted for an approximate length of 26.44 Mb of the 2.13 Gb assigned to homeoblocks, indicating that extreme regulatory divergence between homeoblocks, although overrepresented in certain chromosomes, impacts a minority of the genome.

**Figure 4.**
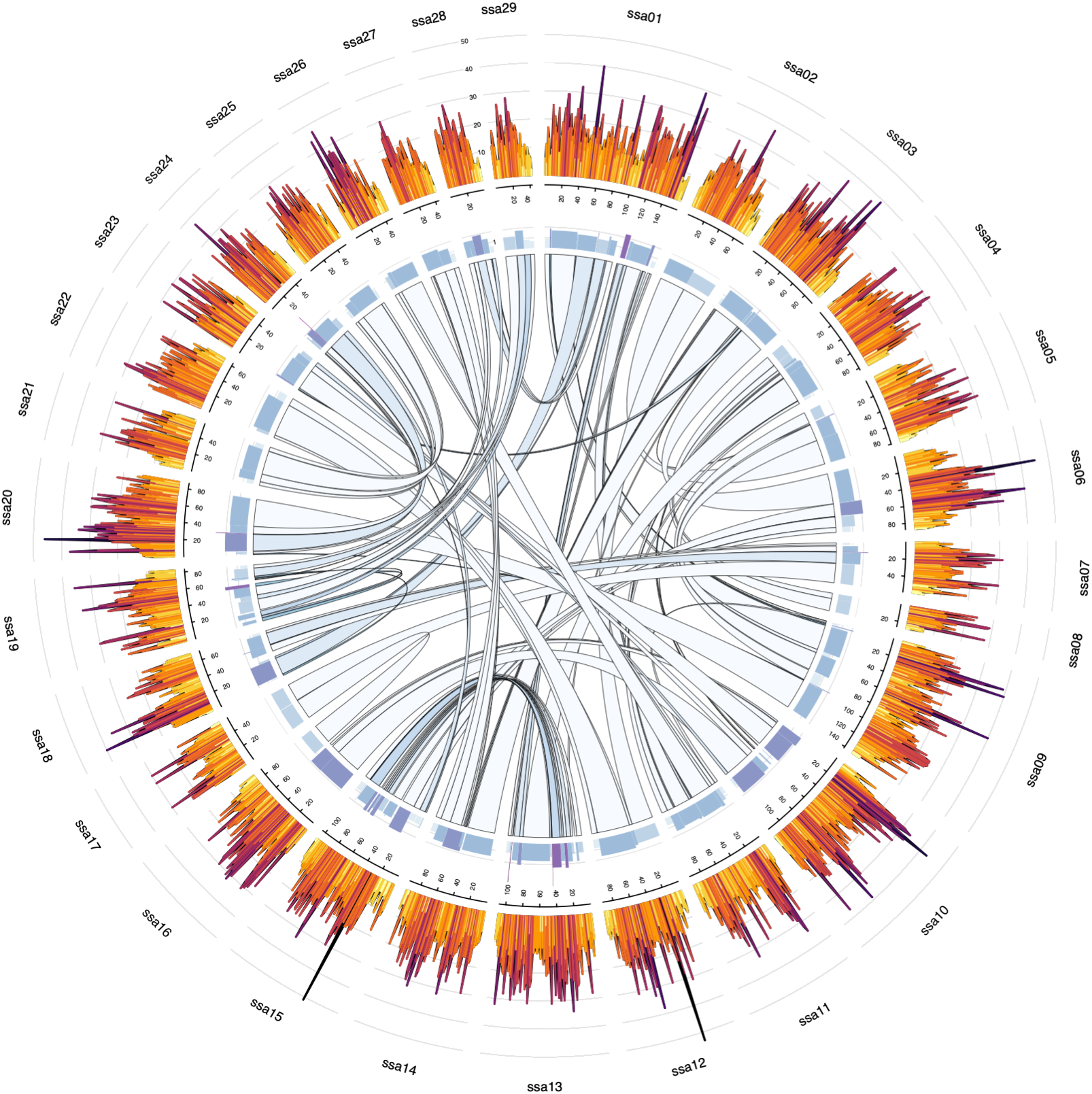
Genomic distribution of H3K27ac peaks. Circos plot showing tracks (from outside) for the number of H3K27ac peaks per 1MB windows, peak density for homeoblocks, and links identifying homeoblocks. Links are colored according to residuals from expected 1:1 peak density between homeoblocks (darker color indicates stronger differences in peak density, similar to Fig 3B).

### Largely neutral evolution of functional elements in natural populations

Our results so far showed that H3K27ac peaks were associated with regulatory activity and divergence in expression levels, indicating that loss or gain of peaks had phenotypic effects in the form of modifying the expression of proximal genes. Yet, ohnologs and homeoblocks tended to show patterns most consistent with random loss or gain in regulatory elements over the timescale of tens of millions of years. To reconcile these observations and to better understand selection on regulatory elements in populations established since the last glaciation 10,000 years ago, we analyzed a population re-sequencing dataset comprised of 31 salmon individuals from across Northern Europe (Barson et al. 2015).

All H3K27ac peaks combined showed comparable Tajima’s *D* and π estimates to putatively neutral 4-fold degenerate sites (Fig 5). When compared to their sequence context, H3K27ac peaks we generally not different from the whole genome. Nucleotide diversity was more strongly reduced in sequence contexts expected to be under stronger purifying selection, with the following order from highest to lowest diversity: introns, intergenic regions, UTRs, CDS regions and 0-fold degenerate sites, respectively (Fig 5). The exception to this were H3K27ac peaks assigned to intronic and intergenic regions that had significantly elevated π (bootstrapping *p*<0.05) compared to the region as a whole. However, these H3K27ac regions did not significantly differ from π at 4-fold degenerate sites. Additionally, Tajima’s *D* at intronic and intergenic H3K27ac peaks was not significantly different from 0, or from *D* at 4-fold degenerate sites.

**Figure 5.**
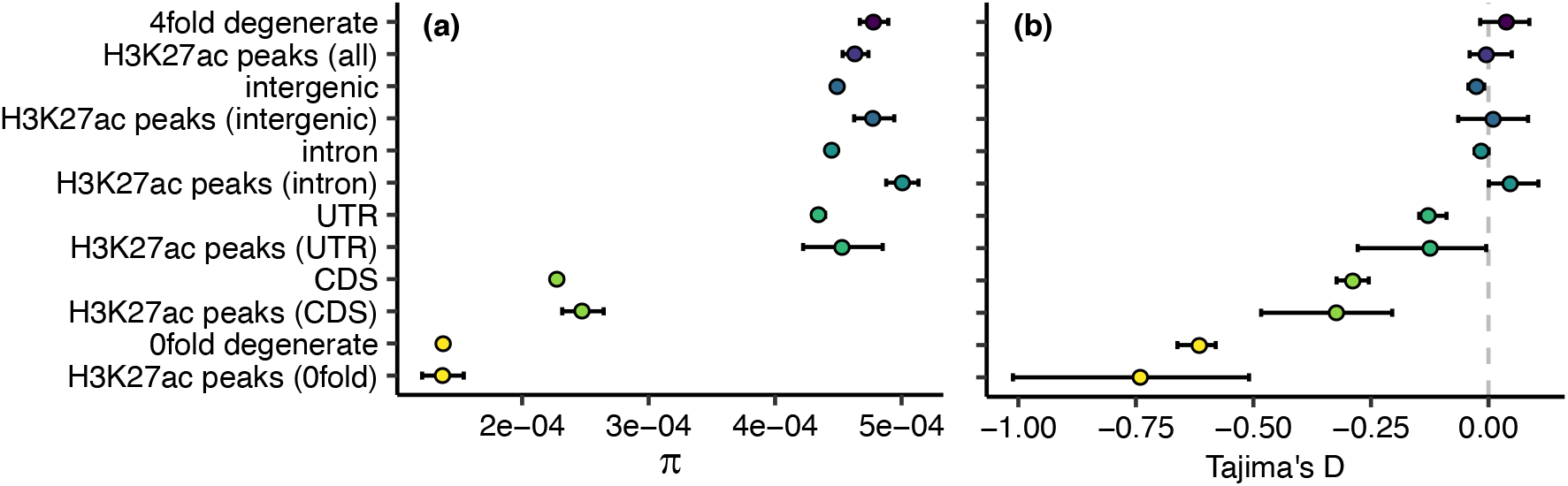
Population genetic summary statistics for H3K27ac peaks reflect their genomic location. (**a**) Nucleotide diversity (π) and (**b**) Tajima’s *D* estimates for 4-fold degenerate (putatively neutral), intergenic, intronic, untranslated region (UTR), coding sequence (CDS), and 0-fold degenerate SNPs, as well as SNPs falling within those regions and within identified H3K27ac peaks, and all SNPs, in all regions, falling within identified peaks. Error bars represent 95% confidence intervals obtained from 100 rounds of bootstrapping by gene.

To further quantify the selective pressures in H3K27ac peaks whilst controlling for the confounding effects of demography, we obtained maximum likelihood estimates of the distribution of fitness effects of new mutations (DFE). For intergenic regions, introns, UTRs, H3K27ac peaks within these regions and all H3K27ac peaks combined we estimated that the majority of point mutations have N_e_s ≤ 1 (Fig 6a), suggesting these regions are evolving neutrally. We additionally used the DFE to calculate the fixation probability of deleterious mutations and consequently the proportion of substitutions fixed by positive selection (α). Slightly negative α values for these regions provided no evidence of positive selection (Fig 6b).

**Figure 6.**
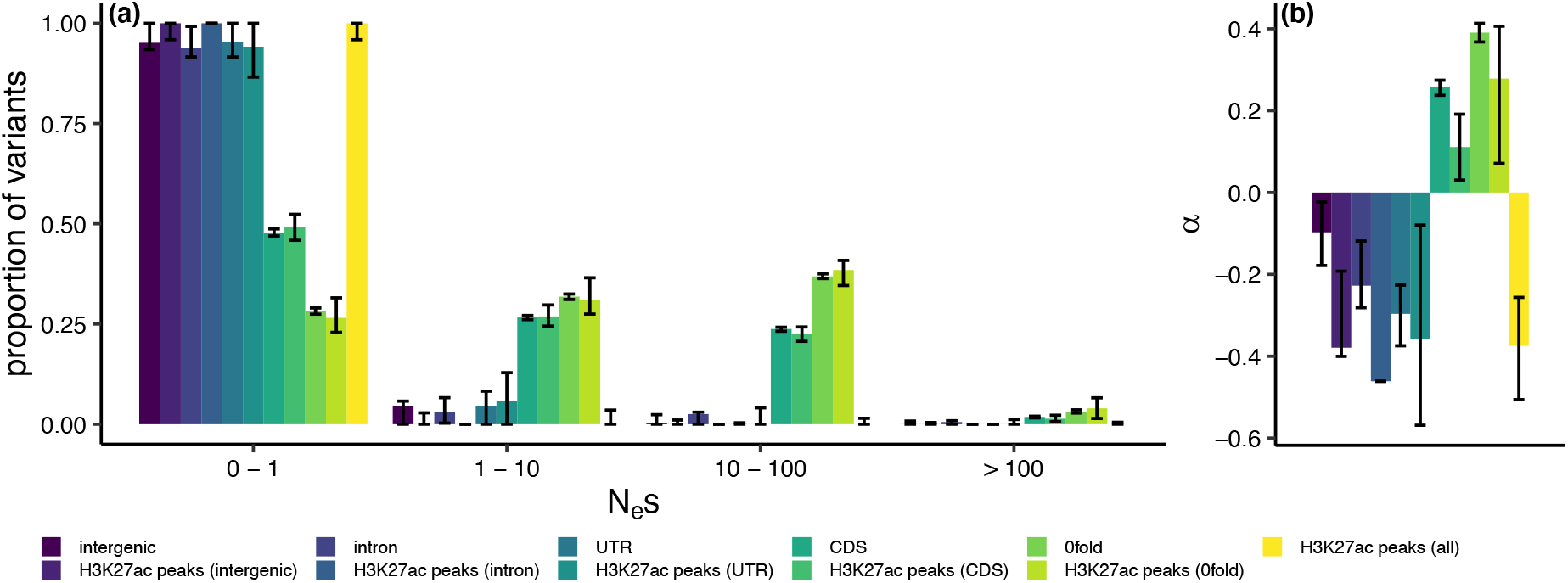
The DFE and α for H3K27ac peaks support predominantly neutral evolution. Estimated distribution of fitness effects of new mutations (**a**; DFE) and the proportion of substitutions fixed by positive selection (**b**; α) in different genomic regions and H3K27ac peaks in those regions, using the population re-sequencing dataset. The DFE is presented as the proportion of variants falling into 4 selective bins: effectively neutral (0 ≤ N_e_s ≤ 1), mildly deleterious (1 < N_e_s ≤ 10), deleterious (10 < N_e_s ≤ 100) and strongly deleterious (N_e_s >100). Error bars represent 95% confidence intervals obtained from 100 rounds of bootstrapping by gene.

The exception to the predominantly neutral pattern of genetic variation were CDS regions and 0-fold degenerate sites, and the H3K27ac peaks within them. These regions were more enriched for SNPs in the weakly deleterious, deleterious and strongly deleterious categories (Fig 6a). Estimates of α in CDS and 0-fold degenerate sites indicated the presence of adaptive substitutions at these sites, not seen in other sequence categories (Fig 6b). We observed a significantly lower α of 0.11 at CDS H3K27ac peaks than at CDS sites as a whole (bootstrapping *p*<0.05), but no significant difference between 0-fold degenerate H3K27ac peak sites and 0-fold degenerate sites in general (Fig 6b).

## Discussion

Molecular evolutionary processes, including the evolution of gene regulatory elements, have remained poorly understood in genomes that contain massive functional redundancy (Elurbe et al. 2017). Analysis of modern genomes derived from whole genome duplications can yield critical insights into regulatory evolution after WGD, specifically pertaining to questions such as the extent and tempo of divergence in regulatory elements between duplicate genome regions, level of regulatory conservation between duplicated genes, and selection forces on regulatory elements. To address these knowledge gaps, we mapped regulatory elements in the Atlantic salmon genome by means of H3K27ac ChIPmentation and analysis of RNA-seq and examined contemporary selection forces on regulatory elements by population resequencing.

Mapping of epigenetic histone modifications through ChIP-seq has been established as a powerful method for inferring putative regulatory elements in metazoan genomes (Andersson and Sandelin 2019). Being dispersed over multiple nucleosomes bordering regulatory elements has in some cases hindered the resolution of histone ChIP-seq signal, and multiple variants of the technique have been developed to yield greater resolution e.g. (Skene and Henikoff 2015). By using a ChIPmentation strategy that implements Tn5-mediated integration of sequencing adapters to a conventional ChIP-seq protocol (Schmidl et al. 2015), our results provide evidence to suggest that acetylated histones can be mapped with near-nucleosome precision in Atlantic salmon using our adapted protocol that is suitable for very small samples. Notably, H3K27ac signal around Atlantic salmon genes showed a distinctive nucleosome patterning with −1, +1, +2 and +3 acetylated nucleosome bordering transcription start sites, as well as distinctive nucleosome free regions immediately upstream of transcription start and downstream of transcription termination sites. These features are a conserved mechanism associated with active transcription (Jiang and Pugh 2009).

We further show an association between H3K27ac peaks and general transcriptional motifs, such as TATA-box, as well as transcription factors such as OCT6 (Pou3f1), SMADs, SF1 and FoxL2. These transcription factors have broadly conserved roles in sexual development. OCT6 is known to be expressed in murine testes and to regulate spermatogonial germ cell renewal (Wu et al. 2010) but to our knowledge has not previously been implicated in fish testis development. SMADs and SF1 are key regulators of testicular cell type differentiation in fish (Sandra and Norma 2010; Pfennig et al. 2015). Notably, the over-representation of SMAD2 and SMAD3 motifs in H3K27ac peaks highlights the importance of the transforming growth factor β (TGF-β) and activin pathways in regulating immature testis differentiation. We also observed an over-representation of FoxL2 motifs, supporting this transcription factor being most often associated with teleost ovary development and may play a role in male gonad development as well (Baron et al. 2005; Liu et al. 2007). Taken together, we anticipate that further assessment of ChIPmentation with additional tissues, epigenetic modifications as well as transcription factors will greatly accelerate the functional annotation of salmonid genomes (Macqueen et al. 2017).

Leveraging on reproducible H3K27ac peaks and publicly available RNA-seq data, we investigated the functional effect that divergence in regulatory elements imparts on gene expression levels between ohnologs. We observed that ohnolog expression divergence was predicted by the difference in the number of proximal peaks assigned to each ohnolog, but not by the difference in promoter peaks. This observation may reflect the lower number of differences in promoter peaks, and hence lower statistical power, but could also have a biological basis in the function of enhancer and promoter elements. Enhancers and promoters evolve at different rates across mammals, where divergence in promoters is notably slower (Villar et al. 2015). Interestingly, (Berthelot et al. 2018) observed a switch-like behavior for promoters, where one promoter element was enough to activate expression and additional promoter elements did not correlate with expression level increase. In contrast, our data show that an increase in promoter peaks correlates with higher expression. Increase in promoter peaks did not however correlate with expression difference between ohnologs, suggesting that divergence in enhancer elements is the primary mechanism for ohnolog expression divergence after WGD.

Genomes from WGD contain duplicated chromosomes that break down into smaller segments, called homeoblocks, over time due to recombination and structural changes. Homeoblocks are known to harbor functional ohnolog genes, yet until now the level and pattern of regulatory element conservation in homeoblocks has remained unclear. By comparing regulatory evolution at the homeoblock level, we observed that differences in H3K27ac peaks normalized to homeoblock length (i.e. peak density) were stronger in shorter homeoblocks. These results fit an interpretation whereby differences in H3K27ac peaks between homeoblocks are largely a product of random loss or gain in peaks, which is predicted to have a larger impact on peak density in shorter homeoblocks. Not all regulatory regions of the Atlantic salmon genome evolve at the same rate, however; retention of polyploid state for a longer period correlated with less regulatory divergence (slower regulatory evolution) between homeoblocks. These results are consistent with lower levels of expression divergence between ohnologs residing in regions with longer polyploid history (Robertson et al. 2017). Longer retained polyploid state also correlated with higher sequence similarity (Lien et al. 2016), which may explain why these regulatory regions show higher conservation in H3K27ac peak densities. It is also worth noting that genes within regions with longer polyploid state corresponded to cases where speciation has preceded ohnolog divergence (Robertson et al. 2017); ohnologs residing in these regions have diverged independently within salmonid species and are therefore good candidates for contributing to species-specific adaptations. Together these results paint a picture which is compatible with massive post-WGD functional redundancy leading to predominantly neutral divergence of regulatory elements between duplicated genome regions, with more restrained evolution in recently polyploid genome regions possibly indicating stronger selection.

Neutral regulatory evolution stemming from functional redundancy is seemingly in contrast with our results that divergence in regulatory elements correlated with phenotypic divergence between ohnologs. Reconciling random regulatory divergence between ohnologs and the functional effects of such evolution requires understanding the selection forces governing genetic diversity in regulatory regions. By genome resequencing and modelling the effects of genetic variation in natural populations, we found that the evolutionary genetic signals of most regulatory elements were indeed consistent with them experiencing largely neutral evolution, not dissimilar to 4-fold neutral sites. The strongest evidence for purifying selection was observed for H3K27ac peaks assigned to UTRs (overlapping promoters and transcription start sites) and coding sequences, while intronic and intergenic H3K27ac peaks, most likely representing enhancer elements, appear evolved largely neutrally. The selection forces acting on these H3K27ac regions however did not differ from their sequence context in general, suggesting that the driving force behind different strengths of selection between H3K27ac peaks was not due to differences in their functionality, but rather their genomic context. Intriguingly, promoter elements show more conserved evolution across mammals as well, compared to enhancer elements (Villar et al. 2015; Berthelot et al. 2018), suggesting that similarly contrasting dynamics between promoter and enhancer elements observed between ohnologs following WGD are also manifested in interspecific regulatory evolution, and likely have similar causes stemming from different genomic contexts. Our results also imply that expression level divergence between ohnologs in the Atlantic salmon genome is presently without major selective effects at large. We anticipate that functional dissection of additional tissues, developmental time points as well as related species will eventually uncover additional evolutionary signatures on regulatory element divergence.

We additionally see that the Atlantic salmon genome has very few strongly deleterious variants (N_e_s > 100) in any genomic region in stark contrast to larger N_e_ species such as many plants (40 – 80% of coding SNPs with N_e_s > 100) (Gossmann et al. 2010), birds (~80% of 0-fold degenerate SNPs with N_e_s > 10 in the great tit and zebra finch)(Corcoran et al. 2017), insects (*Drosophila melanogaster* 78% of nonsynonomous SNPs with N_e_s > 100) (Kousathanas and Keightley 2013), or small mammals (*Mus musculus castaneus* 69% of nonsynonomous SNPs with N_e_s > 100) (Kousathanas and Keightley 2013). This suggests that the salmon N_e_ is currently reduced to the point where selection is unable to act efficiently on many variants, even in coding regions, where over 50% of nonsynonymous mutations seem to be effectively neutral or only weakly deleterious. Thus, any weak selection, such as may be expected at H3K27ac peaks, is likely swamped by genetic drift. However, this is not reflected by our α estimate of ~40% at 0-fold degenerate sites, which represents the proportion of fixations that are adaptive along the whole branch leading to Atlantic salmon from the salmon-brown trout split. Together, these results suggest that from a longer-term perspective, selection has been relatively efficient since the divergence of Atlantic salmon and brown trout.

Studies on gene expression evolution in salmonids support our conclusion that regulatory evolution in salmon follows a predominantly neutral pattern. Tissue expression divergence indicates that asymmetric evolution of expression levels dominates in salmonids (Lien et al. 2016), which is consistent with the largely neutral evolution of gene regulatory elements observed here. Contrasting results showing purifying selection on gene expression levels between salmonids have been reported as well (Varadharajan et al. 2018). Nevertheless, (Gillard et al. 2020) recently found significant evidence for predominant pseudogenization of expression levels (neutral loss of expression patterns) across salmonids. We have shown here that the most likely explanation for such predominant pattern of pseudogenization of ohnolog expression patterns is likely coupled with largely neutral *cis*-regulatory evolution. Predominantly neutral evolution of regulatory sequences may also help explain the observed prevalence of pseudogenization of ohnologs in the absence of strong coding sequence divergence (Lien et al. 2016).

Our study demonstrates that an ancient WGD continues to have an impact on the regulatory landscape of the Atlantic salmon genome after at least 80 million years of evolution. Overall, massive functional redundancy could potentially lift selective constraints on most gene regulatory elements, with a notable exception of gene promoters and coding sequences, which among regulatory elements showed the strongest signals of past selection. Genome-wide, the tempo of regulatory evolution has followed a varied, with parts of the genome that experienced longer polyploidy showing less regulatory divergence. We anticipate that future studies will uncover selected regulatory elements within this largely neutrally evolving post-WGD regulatory genome.

## Acknowledgements

This work was supported by European Research Council (grant agreement number 742312), Academy of Finland (grant numbers 314254 and 314255) and the university of Helsinki. We thank the Primmer lab members for help in fish husbandry and sampling, the LUKE staff with initial crosses and Kai Zeng for providing example code to calculate alpha.

## Material and Methods

### Experimental design and collection of material

We collected immature male gonads from five 11-12-month-old male Atlantic salmon raised in common-garden conditions (see details in (Verta et al. 2020). Fish were euthanized using an overdose of MS-222, followed by decapitation. Gonads were dissected under a microscope, flash-frozen in liquid nitrogen and stored in −80 degrees C until use for chromatin extraction.

### H3K27ac ChIPmentation

Detailed protocol for ChIPmentation (Schmidl et al. 2015) of salmon gonad tissues is provided in Supplementary file X. Briefly, we integrated ChIPmentation to the workflow from ThermoFisher MAGnify ChIP kit. Gonads were homogenized in D-PBS buffer using OMNI Beadruptor Elite device in 2 ml tubes and 2.8mm stainless steel beads. Chromatin was fixed using 1% formaldehyde for 2 min, followed by quenching using 0.125 M glycerine concentration for 5 min. Cells were collected using centrifugation and resuspended in lysis buffer supplemented with protease inhibitors. Chromatin was sheared in 150 ul volumes using a Bioruptor device with settings high power and 8 cycles of 30 sec on, 30 sec off. Debris was pelleted by centrifugation and sheared chromatin was diluted to IP conditions. An aliquot of sheared chromatin was reserved as input control. Acetylated histones were immunoprecipitated in +4 degrees C for 2 hours using 1 microgram of SantaCruz ab4729 on ThermoFisher Dynabeads Protein A/G. Beads were subsequently washed following MAGnify kit protocol, with an additional final wash using 10 mM Tris (pH 8). Bead-bound chromatin was then treated with a tagmentation reaction containing Illumina Tn5 transposase for 5 min at 37 degrees C. Tagmentation was terminated by adding 7.5 volumes of RIPA buffer and incubation on ice for 5 min. ChIPmented chromatin was subsequently washed twice with both RIPA and TE buffer. Crosslinks were reversed using a proteinase-K treatment and ChIPment DNA was captured using magnetic beads. These steps were performed following the MAGnify kit protocol (for 3 samples), or alternatively by using a reverse crosslinking buffer (10 mM Tris-Hcl pH8, 0.5% SDS, 300 mM NaCl, 5 mM EDTA, proteinase-K) and Macherey-Nagel NucleoMag magnetic beads (for 2 samples). Input controls were treated with tagmentation reaction for 5 min at 55 degrees C. Tn5 was inactivated by adding SDS and tagment DNA was purified using Macherey-Nagel NucleoMag magnetic beads. Successful adapter integration was tested using PCR and primers aligning with Nextera adapters.

### Alignment of ChIP-seq reads

ChIPmentation and matched input control libraries were sequenced using Illumina Nextseq chemistry at the Institute of Biotechnology of the university of Helsinki. Libraries were sequenced using both single-end and paired-end strategies, and the resulting ChIP fragment directories combined for each sample as described in the following section. For single-end libraries, reads were passed through a quality-control including Nextera adapter trimming using *fastp* (Chen et al. 2018) and the following parameters *--low_complexity_filter --trim_tail1=1 --trim_front1=19.* Reads were then aligned to the Atlantic salmon genome (Lien et al. 2016) downloaded from NCBI (version: ICSASG_v2, available from: ftp://ftp.ncbi.nlm.nih.gov/genomes/all/GCF/000/233/375/GCF_000233375.1_ICSASG_v2/GCF_000233375.1_ICSASG_v2_genomic.fna.gz) using *bowtie2 v2.4.2* (Langmead and Salzberg 2012) and the following parameters *--very-sensitive --end-to-end*. Paired-end libraries were correspondingly analysed using *fastp* (*--low_complexity_filter --trim_front1=19 --trim_tail1=50 --trim_tail2=12*) and *bowtie2* (*--very-sensitive --maxins 1500 --end-to-end*). Alignment files were then quality filtered using *samtools* (Li et al. 2009) and parameters *-F 256 -q 20*.

### Identification and annotation of H3K27ac peaks

We used *HOMER* (Heinz et al. 2010) to call enriched H3K27ac regions over input control and to annotate the reproducible peaks. ChIP fragment distributions were created using the command *makeTagDirectory* and parameters -*keepOne -single -tbp 1 -mis 5 -GCnorm default*. Tag directories for single-end and paired-end libraries for the same samples were then combined similarly with a *makeTagDirectory* command. Reproducible H3K27ac regions were identified with the command *getDifferentialPeaksReplicates.pl*, specifying parameters *-style histone.* Results were transformed into a bed file with *pos2bed.pl*. We then used custom *R* (Team 2013) code and *bedtools intersect* (Quinlan and Hall 2010) to filter the peaks for any overlapping 1 Kb genomic windows with the top 1% of mean sequencing coverage to avoid problematic regions of the genome. Peaks were then annotated using a *annotatePeaks.pl* command and parameters *-CpG* and *HOMER* motif search was performed using *findMotifsGenome.pl* and parameters *-size given*. Finally, specific instances of motifs and their coverage were identified using *annotatePeaks.pl* and parameters *-m*, and parameters *-size 4000 -hist 10 -m -d*.

### RNA-seq alignment and quantification

RNA-seq reads of immature male gonads (SRR8479243, SRR8479245, SRR8479246) (Skaftnesmo et al. 2017) as well as 14 salmon tissues from (Lien et al. 2016) were downloaded from *Sequence Read Archive* and filtered using *fastp* and default parameters. We used *STAR* (Dobin et al. 2013) to create a genome index (-*runMode genomeGenerate*) and align RNA-seq reads to the Atlantic salmon genome downloaded from *NCBI,* in manual two-pass mode, with the following parameters *— outFilterIntronMotifs RemoveNoncanonicalUnannotated -chimSegmentMin 10 – outFilterType BySJout -alignSJDBoverhangMin 1 -alignIntronMin 20 -alignIntronMax 1000000 -alignMatesGapMax 1000000 -quantMode GeneCounts – alignEndsProtrude 10 ConcordantPair —limitOutSJcollapsed 5000000*. Alignments were quantified over gene models downloaded from *NCBI* using *R* function *featurecounts* from the *Rsubread* package (Liao et al. 2019) and normalized using *DESeq2 varianceStabilizingNormalization* (Love et al. 2014) (immature male gonads) or RPKM (14 tissues).

### Analysis of population resequencing data

#### Read Mapping

We downloaded the raw reads for the whole genome resequencing dataset described in (Barson et al. 2015) from the European Nucleotide Archive (ENA) under study accession PRJEB10744 (available from: https://www.ebi.ac.uk/ena/browser/view/PRJEB10744). The dataset comprises 31 salmon individuals from seven populations across the Atlantic and Barents Sea sequenced using the Illumina HiSeq 2500 platform (125 bp, paired end reads).

We cleaned the reads and removed adapter contamination using *Trim Galore* (version: 0.6.4_dev) (available from: https://github.com/FelixKrueger/TrimGalore) with *Cutadapt* (version: 2.7) (Martin 2011). We then aligned the reads to Atlantic salmon reference genome using *BWA-MEM* (version: 0.7.17-r1188) (Li 2013) and marked PCR duplicates and added read group information using *Picard* (version: 2.22.4) (Institute n.d.). This resulted in a mean mapped coverage of 8X (see table S1 for individual sample coverages).

#### Variant Calling

We followed the GATK (version: 4.1.3.0) pipeline to call SNPs (Auwera et al. 2013), and restricted our dataset to assembled chromosomes only. First, we generated a training set for use in base quality score recalibration (BQSR) and variant quality score recalibration (VQSR) by intersecting an initial SNP call set obtained using *HaplotypeCaller* and *GenotypeGVCF* in GATK with a call set obtained using *SAMtools* (version: 1.9). We further subset the training set by intersecting it with the positions of variants used on the 20k and 200k salmon SNP chips. Second, we performed BQSR using this training set to produce recalibrated BAM files. Third, we called SNPs from the recalibrated BAM files using *HaplotypeCaller* and *GenotypeGVCF* in GATK. Forth we performed VQSR, retaining variants that passed a tranche level cut-off of 99.5%. Finally, we filtered out SNPs that were multi-allelic, fell in repetitive regions, had a mean depth below 4X or above 16X (half and twice the mean coverage) and that were missing genotype calls in some individuals. This resulted in a dataset of 3,723,849 SNPs.

### Whole Genome Alignment and Ancestral States

To infer the ancestral alleles of the SNPs in our dataset we used a 9-way multispecies whole genome alignment and maximum parsimony using the brown trout (*Salmo trutta*) Atlantic salmon (*Salmo salar*), Arctic charr (*Salvelinus alpinus*) sequences; at each bi-allelic SNP in salmon, to assign an allele as ancestral we required it matched the sequence in the other two species. This resulted ancestral alleles inferred for 1,614,400 out of 3,723,849 SNPs.

The whole genome alignment was performed as follows. Pairwise alignments were generated between each brown trout chromosome and each query species genome (a full list of species and genome versions can be seen in table SX) using *LASTZ* (Harris and R.S. 2007). The pairwise chromosomal alignments were then chained using *axtChain* and netted with *chainNet* (Kent et al. 2003) The chromosomal alignments were merged into whole genome alignments and single coverage was insured for the reference sequence (the brown trout reference genome) using *single_cov2.v11* from the *MULTIZ* package (Blanchette et al. 2004). *MULTIZ* was then used to align the pairwise alignments using the automation script *roast*.

### Annotating Variants

We identified SNPs falling in different genomic regions (intergenic, intronic, untranslated regions [UTRs], coding sequence [CDS]) by creating bed files of their coordinates from the NCBI GFF file for version CSASG_v2 of the Atlantic salmon genome (available from: ftp://ftp.ncbi.nlm.nih.gov/genomes/all/GCF/000/233/375/GCF_000233375.1_ICSASG_v2/GCF_000233375.1_ICSASG_v2_genomic.gff.gz). To obtain genomic coordinates for fourfold degenerate sites we parsed the CDS fasta file for the same genomic version (available from: ftp://ftp.ncbi.nlm.nih.gov/genomes/all/GCF/000/233/375/GCF_000233375.1_ICSASG_v2/GCF_000233375.1_ICSASG_v2_cds_from_genomic.fna.gz) with a custom python script. Finally we obtained coordinates for repeat regions in the genome from the repeat masker output from NCBI (available from: ftp://ftp.ncbi.nlm.nih.gov/genomes/all/GCF/000/233/375/GCF_000233375.1_ICSASG_v2/GCF_000233375.1_ICSASG_v2_rm.out.gz).

### Summary Statistics

We calculated nucleotide diversity (*μ* (Tajima 1983)) and Tajima’s *D* (Tajima 1989) for CDS sites, 4-fold degenerate sites, UTRs, introns as well as sites within enhancers, subdivided by their genomic location (intergenic, intronic and UTRs). To extract variants within these regions we intersected our VCF file with the relevant BED file generated from our annotation pipeline, using *BEDTools* (version: 2.26.0).

To obtain per site estimates of *μ* we created a custom FASTA file containing all sites in the genome that passed our variant calling filters (see Variant Calling above), by applying the filters to an all sites (monomorphic sites and variants) VCF file output during variant calling, from GATK’s *HaplotypeCaller* with the *-ERC GVCF* flag. From this file we calculated the number of callable sites for each genomic region.

### Distribution of Fitness Effects

We estimated the distribution of fitness effects (DFE) using the package *anavar* (version 1.2) (Barton and Zeng 2018). Briefly, the method uses the site frequency spectrum to estimate the population scaled mutation rate (*θ* = 4N_e_*μ*) and shape and scale parameters for a gamma distribution of population scaled selection coefficients (γ = 4N_e_s). Here we present the gamma distributions as the proportion of variants falling into 4 selective bins, effectively neutral (0 ≤ N_e_s ≤ 1), mildly deleterious (1 < N_e_s ≤ 10), deleterious (10 < N_e_s ≤ 100) and strongly deleterious (N_e_s >100).

We fitted the *neutralSNP_vs_selectedSNP* model, with a continuous gamma distribution to model the strength of selection, separately to CDS sites, UTRs, introns and sites within enhancers, subdivided by their genomic location (intergenic, intronic and UTRs). The model takes the unfolded site frequency spectrum for a focal set of sites and estimates the population scaled mutation rate (*θ* = *4N*_*e*_*μ*, where N_e_ is the effective population size and μ is the per site per generation mutation rate), the shape and scale parameters for a gamma distribution of population scaled selection coefficients (γ = *4N*_*e*_*s*, where s is the selection coefficient) and the rate of ancestral state misidentification (polarization error; *ε*), which is used internally to correct the SFS. The model also uses the site frequency spectrum from a putatively neutral set of sites to control for the confounding effects of demography following the method of (Eyre-Walker et al. 2006). Here we used the SFS from fourfold degenerate sites for this purpose. We present our DFE results by using the estimated gamma distribution of selection coefficients to estimate the proportion of variants falling into four γ bins (note all γ estimates are negative), effectively neutral (0 ≤ γ ≤ 1), weakly deleterious (1 < γ ≤ 10), deleterious (10 < γ ≤ 100) and strongly deleterious (γ >100).

### Bootstrapping

To obtain 95% confidence intervals for our summary statistics and DFE analyses we performed 100 rounds of resampling with replacement by gene for each SFS dataset. That is, for a given focal SFS we resampled both the focal SFS and the neutral reference SFS (4-fold degenerate sites) and the number of callable sites for each per gene. We then recalculated π and Tajima’s D, and re-estimated the DFE for each bootstrap replicate.

### Calculating α

In order to calculate the proportion of substitutions fixed by positive selection (α), we first obtained divergence estimates for each genomic region in the DFE analyses and for 4-fold degenerate sites. To that end, we created concatenated FASTA alignments for each region, using the Atlantic salmon, Arctic charr and brown trout sequences in our whole genome alignment. We then used the APE package (version 5.4.1) (Paradis et al. 2004) in R (version 3.5.1) to estimate the pairwise distance matrix between species using *dist.dna* with the *K80* model, which we then used to calculate divergence on the Atlantic salmon branch since its split from brown trout.

Second, we estimated the fixation probabilities for deleterious mutations 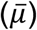 in each genomic region using our DFE estimates with:

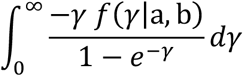

where *f*(γ|a, b) is the probability density function of the reflected Γ distribution of fitness effects, with a the shape parameter and b the scale parameter, as estimated by *anavar*.

Finally, we substituted 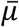 into (equation 19 from (Barton and Zeng 2018)):

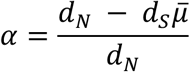

Where d_N_ is our divergence estimate for our focal sites from the DFE analysis and dS is the divergence estimate for 4-fold degenerate sites (our neutral reference).

### Code availability and command lines

For the re-sequencing data preparation and subsequent analysis all scripts and command lines are available on GitHub (available from: https://github.com/henryjuho/sal_enhancers).

### Downstream analyses

We identified ohnologs using best reciprocal BLASTp matches and corresponding homeoblock localization (Lien et al. 2016) using a custom *Python 3* script. All subsequent analyses were performed with custom scripts in *R*, available at the Dryad repository (doi: XXX).

---

**Figure S1.**
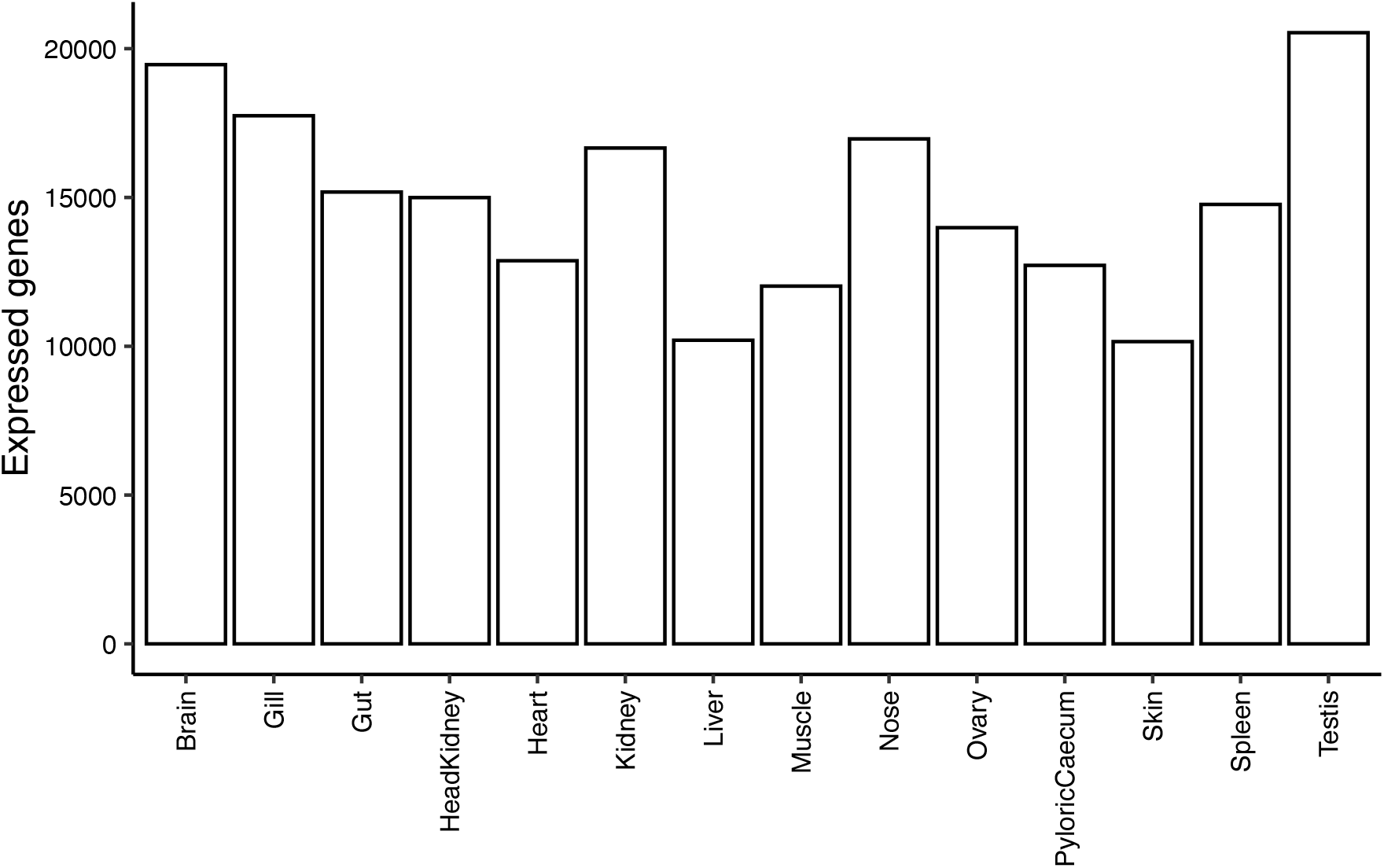
Number of expressed genes (RPKM > 1) in 14 salmon tissues from (Lien et al. 2016).

**Figure S2.**
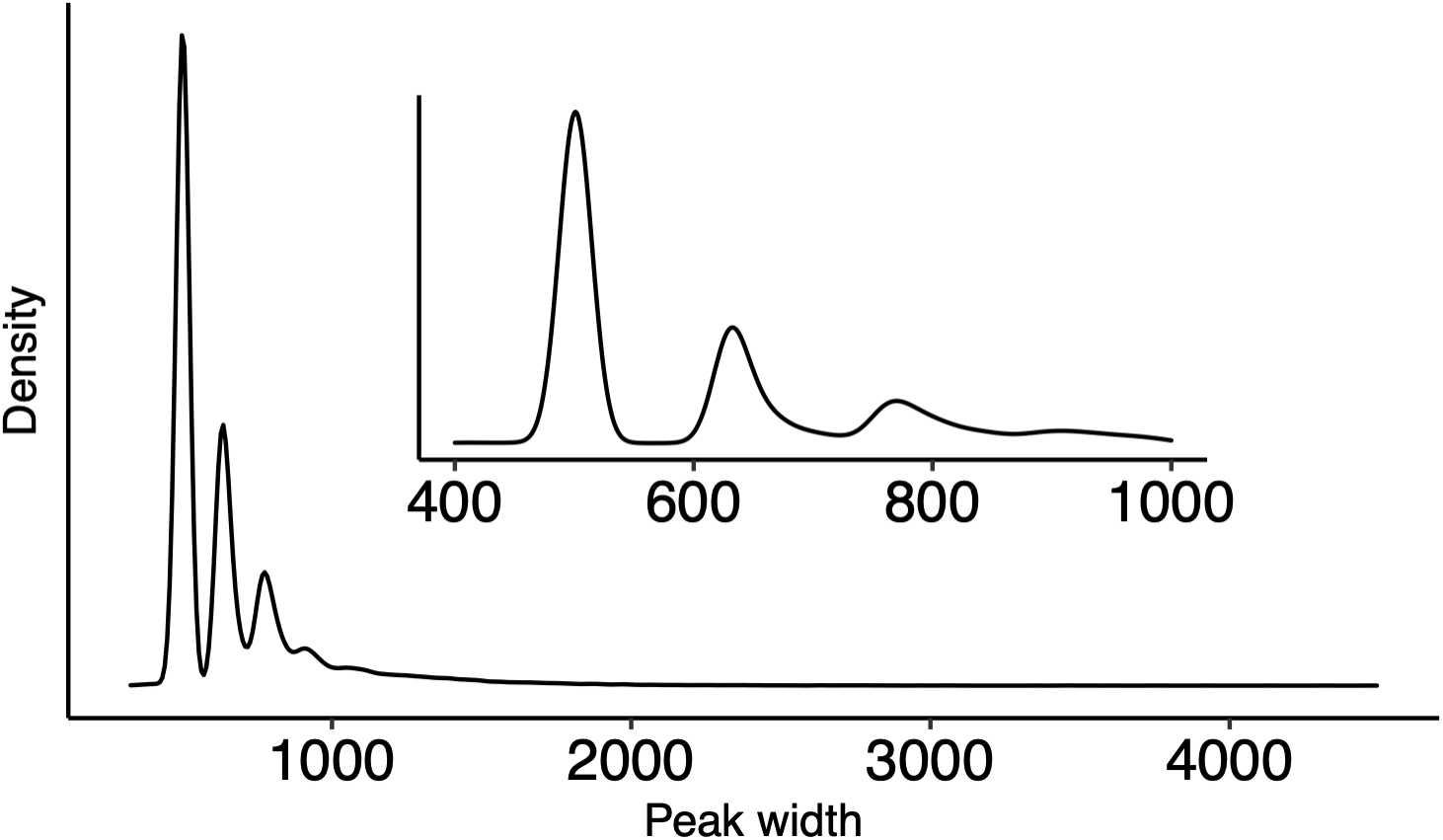
Distribution of H3K27ac peak widths identified using *HOMER*. The default peak width of 500 bp was used. Distribution of peak widths shows a periodical pattern that is consistent with peaks representing regions covering different numbers of acetylated nucleosomes.

**Figure S3.**
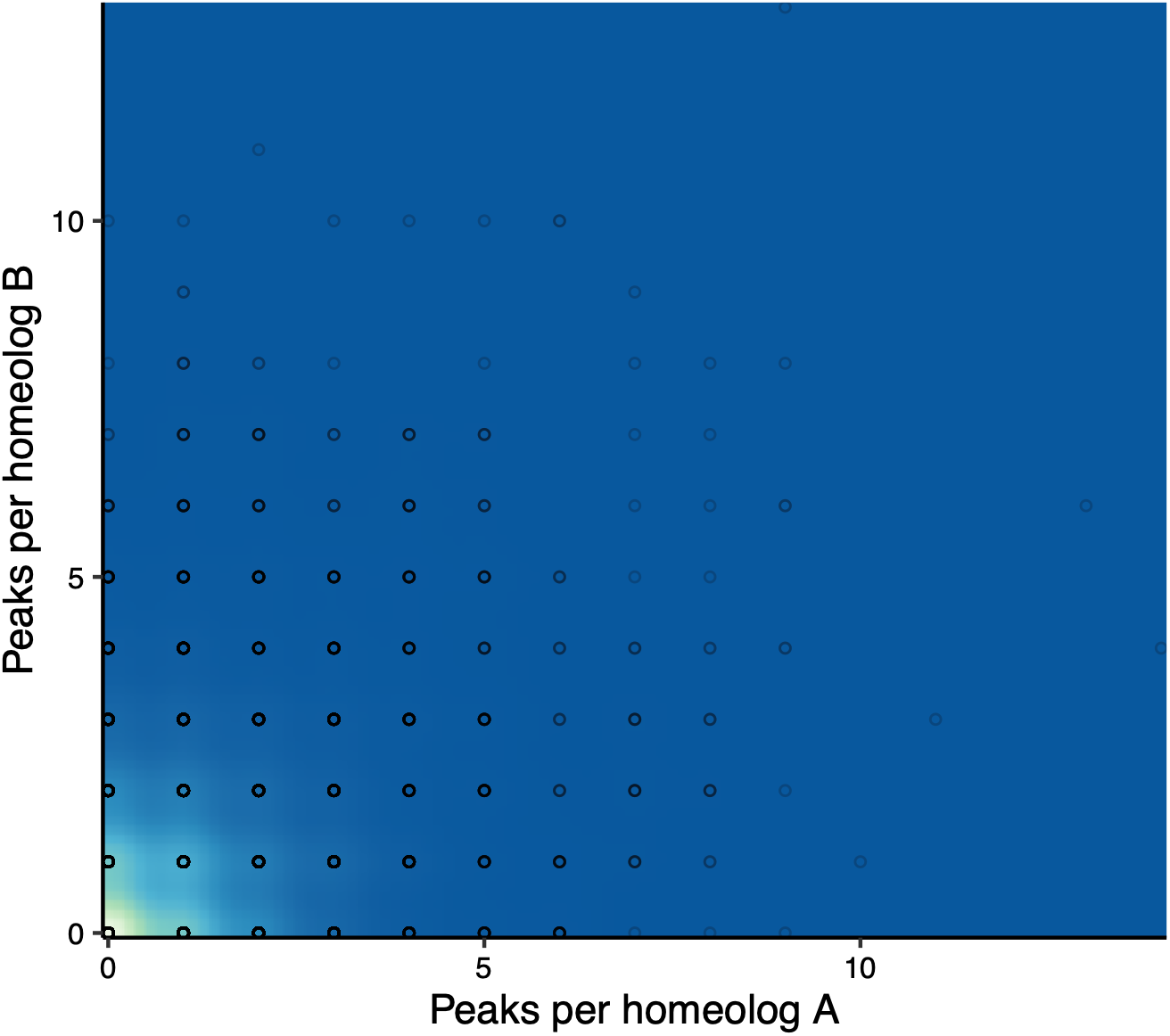
The number of H3K27ac peaks assigned to ohnologs. Background is coloured according to the density of datapoints.

**Figure S4.**
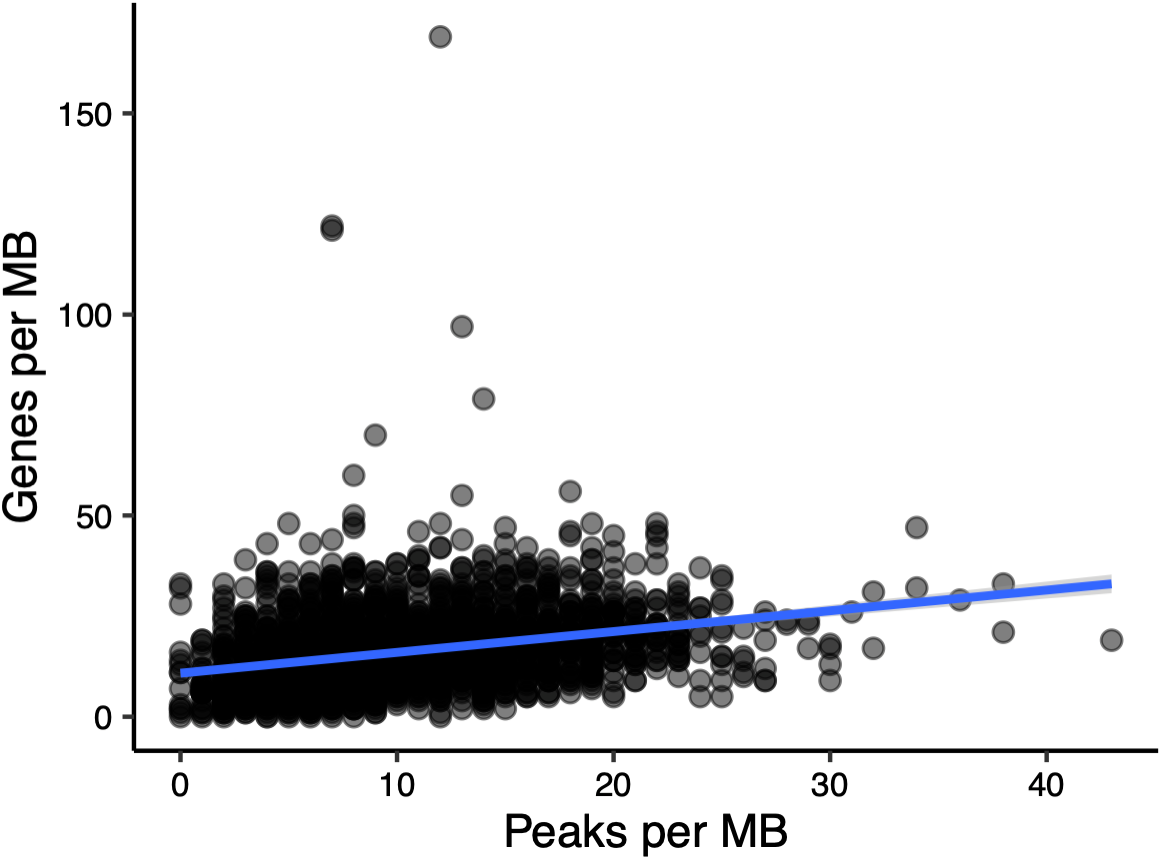
Density of genes versus density of H3K27ac peaks per 1Mb windows.

**Table S1.** Accession ID, sample ID and mean and median sequencing coverage for each sample in the re-sequencing dataset. Measured after mapping the raw reads from Barson et al. (2015) to the Atlantic salmon reference genome and removing PCR duplicates, prior to variant calling.

| run accession | sample | mean coverage | median coverage |
| --- | --- | --- | --- |
| ERR1013407 | Alta_12_0001 | 7.520952 | 8 |
| ERR1013495 | Alta_12_0124 | 3.349194 | 3 |
| ERR1013583 | Alta_12_0228 | 7.705269 | 8 |
| ERR1013463 | Arga_12_0075 | 7.728542 | 8 |
| ERR1013471 | Arga_12_0082 | 9.305405 | 10 |
| ERR1013479 | Arga_12_0089 | 13.245793 | 14 |
| ERR1013647 | Jols_13_0001 | 10.631584 | 11 |
| ERR1013639 | Jols_13_0009 | 8.723167 | 9 |
| ERR1013655 | Jols_13_0027 | 7.64258 | 8 |
| ERR1013447 | Nams_12_0024 | 10.828244 | 11 |
| ERR1013455 | Nams_12_0071 | 10.654392 | 11 |
| ERR1013439 | Nams_12_0201 | 6.65685 | 7 |
| ERR1013423 | Naus_12_0014 | 9.373985 | 10 |
| ERR1013415 | Naus_12_0037 | 7.585018 | 8 |
| ERR1013431 | Naus_12_0059 | 12.050208 | 13 |
| ERR1013615 | Repp_12_0007 | 8.360628 | 9 |
| ERR1013623 | Repp_12_0023 | 6.412691 | 6 |
| ERR1013631 | Repp_12_0034 | 12.499478 | 13 |
| ERR1013487 | Uts_11_15 | 7.155256 | 7 |
| ERR1013511 | Uts_11_17 | 19.169955 | 21 |
| ERR1013519 | Uts_11_24 | 9.167601 | 10 |
| ERR1013527 | Uts_11_26 | 5.419665 | 5 |
| ERR1013535 | Uts_11_27 | 6.556109 | 7 |
| ERR1013543 | Uts_11_28 | 6.834338 | 7 |
| ERR1013551 | Uts_11_29 | 4.531277 | 4 |
| ERR1013559 | Uts_11_30 | 6.543761 | 7 |
| ERR1013567 | Uts_11_31 | 6.588743 | 7 |
| ERR1013575 | Uts_11_39 | 2.994131 | 3 |
| ERR1013591 | Uts_11_46 | 7.094451 | 7 |
| ERR1013599 | Uts_11_52 | 3.360987 | 3 |
| ERR1013607 | Uts_11_53 | 8.927331 | 9 |

**Table S2.**
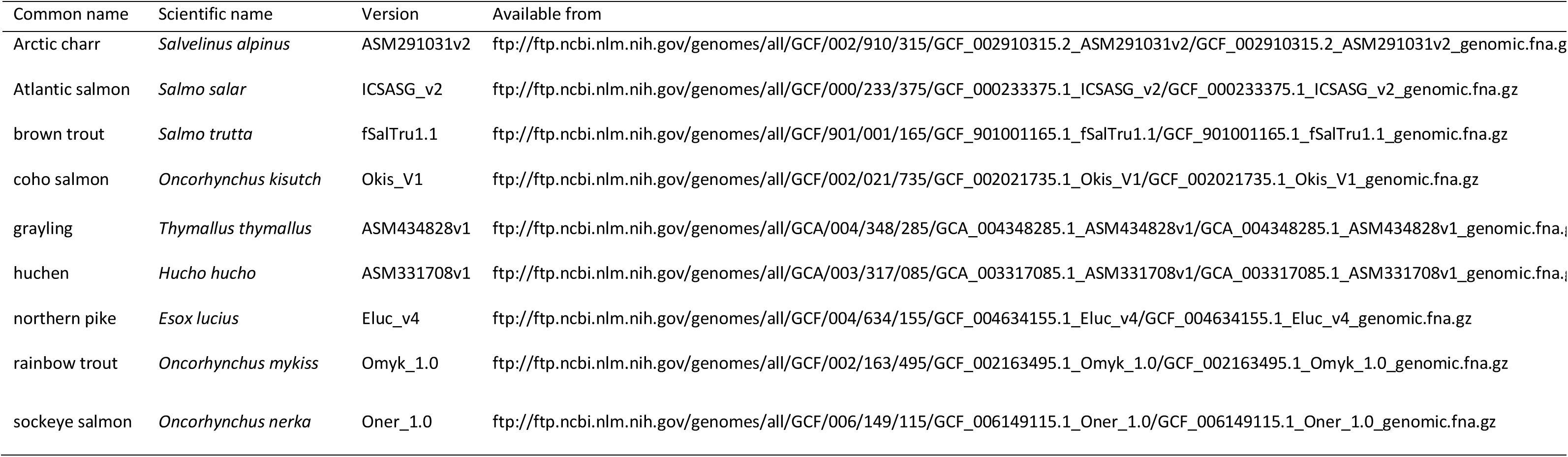
Reference versions and ftp links for genomes used in the multispecies whole genome alignment. Reference species in bold (brown trout)

## Supplementary Analyses

We used alternative parametrization of *HOMER* to H3K27ac peaks that were suitable for transcription factor motif finding (fixed 500 bp peaks centered on nucleosome free regions). This analysis identified 26,507 replicated peaks, of which 83% were overlapping peaks identified in the main analysis. We additionally identified TF motifs in the original peak set, which gave congruent results. Replicated peaks were over-represented in motifs for general promoter features (TATA-box 28.4% of peaks, *p*=1e-196), as well as for transcription factors implicated in the development of spermatogonia (OCT6 10.1% of peaks, *p*=1e-294), Sertoli and Leydig cells (SMAD4 49.3% of peaks *p*=1e-90, SMAD2 47.3% of peaks *p*=1e-51, SF1 12.2% of peaks *p*=1e-132 and FoxL2 41.9% of peaks *p*=1e-75).

We defined a set of one-to-one ohnologs using an alternative approach as implemented in the software *OrthoFinder* (Emms and Kelly 2015) to test whether the results were robust to analysis strategies. We used northern pike (*Esox lucius*) and rainbow trout (*Oncorhynchus mykiss*) as outgroups in the analysis following *OrthoFinder* basic procedure. Results using this alternative ohnolog set did not differ from the original analysis.

